# Antifungal activity of fibrate-based compounds and substituted pyrroles inhibiting the enzyme 3-hydroxy-methyl-glutaryl-CoA reductase of *Candida glabrata* (CgHMGR), and decreasing yeast viability and ergosterol synthesis

**DOI:** 10.1101/2021.09.14.460412

**Authors:** Damián A. Madrigal-Aguilar, Adilene Gonzalez-Silva, Blanca Rosales-Acosta, Celia Bautista-Crescencio, Jossué Ortiz-Álvarez, Carlos H. Escalante, Jaime Sánchez-Navarrete, César Hernández-Rodríguez, Germán Chamorro-Cevallos, Joaquín Tamariz, Lourdes Villa-Tanaca

**Affiliations:** Departamento de Química Orgánica, Escuela Nacional de Ciencias Biológicas, Instituto Politécnico Nacional, Prol. Carpio y Plan de Ayala S/N, CP 11340, Mexico City, Mexico., MX; Departamento de Microbiología, Escuela Nacional de Ciencias Biológicas, Instituto Politécnico Nacional, Prol. Carpio y Plan de Ayala S/N, CP 11340, Mexico City, Mexico; Laboratorio de Investigación Microbiológica, Hospital Juárez de México, Av. Instituto Politécnico Nacional Núm. 5160, México City, Mexico; Departamento de Farmacia, Escuela Nacional de Ciencias Biológicas, Instituto Politécnico Nacional, Prol. Carpio y Plan de Ayala S/N, CP 11340, México City, Mexico

**Author notes:** Corresponding authors: Lourdes Villa-Tanaca.; Laboratorio de Biología Molecular de Bacterias y Levaduras, Departamento de Microbiología, Escuela Nacional de Ciencias Biológicas, Instituto Politécnico Nacional, Prol. de Carpio y Plan de Ayala. Col. Sto. Tomás, 11340 Mexico City, Mexico., Joaquín Tamariz. Departamento de Química Orgánica, Escuela Nacional de Ciencias Biológicas, Instituto Politécnico Nacional, Prol. de Carpio y Plan de Ayala. Col. Sto. Tomás, 11340 Mexico City, Mexico. Both authors contributed to the same extent (Damián A. Madrigal-Aguilar, Adilene Gonzalez-Silva).

**Keywords:** HMGR, ergosterol, fibrates, pyrroles, atorvastatin, synthetic antifungal, *Candida*, multi-drug resistance

## Abstract

Due to the emergence of multi-drug resistant strains of yeasts belonging to the *Candida* genus, there is an urgent need to discover antifungal agents directed at alternative molecular targets. The aim of the current study was to evaluate the capacity of synthetic compounds to inhibit the *Candida glabrata* enzyme denominated 3-hydroxy-methyl-glutaryl-CoA reductase (CgHMGR), and thus affect ergosterol synthesis and yeast viability. One series of synthetic antifungal compounds were analogues to fibrates, a second series had substituted 1,2-dihydroquinolines and the third series included substituted pyrroles. α-asarone-related compounds **1c** and **5b** with a pyrrolic core were selected as the best antifungal candidates. Both inhibited the growth of fluconazole-resistant *C. glabrata* 43 and fluconazole-susceptible *C. glabrata* CBS 138. A yeast growth rescue experiment based on the addition of exogenous ergosterol showed that the compounds act by inhibiting the mevalonate synthesis pathway. A greater recovery of yeast growth occurred for the *C. glabrata* 43 strain and after the **1c** (versus **5b**) treatment. Given that the compounds decreased the ergosterol concentration in the yeast strains, they probably target the ergosterol synthesis. According to the docking analysis, the inhibitory effect of the **1c** and **5b** could possibly be mediated by their interaction with the amino acid residues of the catalytic site of CgHMGR. Since **1c** displayed higher binding energy than α-asarone and **5b**, it is a good candidate for further research, which should include structural modifications to increase its specificity and potency as well as *in vivo* studies on its effectiveness at a therapeutic dose.

**HIGHLIGHTS:**

1. Fibrate-based and pyrrole-containing compounds were tested as *C. glabrata* inhibitors.
2. The best inhibitor from fibrate was **1c** and from pyrroles was **5b**.
3. These agents inhibited *C. glabrata* growth better than the reference antifungals.
4. They also inhibited ergosterol synthesis by the two *C. glabrata* strains tested. Experimental

## INTRODUCTION

The emergence of multi-drug resistant strains of *Candida* yeasts in recent years has made infections by these pathogens a more serious problem [1]. Although *Candida albicans (C. albicans), C. glabrata, C. tropicalis, C. parapsilosis* and *C. krusei* are species isolated from healthy individuals, they can behave as invasive opportunistic pathogens under host conditions of a compromised immune system.

Among the particularly important *Candida* species with multi-drug resistance are *C. auris*, the species of the *C. haemulonii* complex, and *C. glabrata*. The former two cause in-hospital outbreaks and polymicrobial infections associated with SARS-Cov-2 [2,3]. *C. glabrata* is intrinsically resistant to azoles, and the recent pan-echinocandin-resistant strains of this species are also associated with the COVID-19 pandemic [4]. *C. glabrata* has been proposed as a model for the study of statins as antifungal agents [5].

Three main mechanisms of antifungal action have been found to date for antifungal agents: an alteration of the fungal membrane by the binding of polyenes to ergosterol, of the synthesis of ergosterol by the activity of azoles, allylamines and thiocarbamates, and of the generation of the cell wall by echinocandins [6]. A possible alternative target is 3-hydroxy-methyl-glutaryl-CoA (HMGR), an enzyme that catalyzes the synthesis of mevalonate, one of the critical steps in the ergosterol biosynthesis pathway [7-10]. The purpose of developing new antifungals with alternative molecular targets is to provide a wide range of compounds to respond to the multi-drug resistance of *Candida* spp. and other fungi.

The aim of the present study was to evaluate the capacity of new synthetic compounds to inhibit the *C. glabrata* HMGR enzyme (CgHMGR) and therefore affect ergosterol synthesis and yeast viability. Two series of compounds were derived from fibrate-based acyl- and alkyl-phenoxyacetic methyl esters, and 1,2-dihydroquinolines [11] and a third series from substituted pyrroles [12,13]. The best compound in each series was subjected to *in vitro* experiments to assess yeast growth, the level of ergosterol, and yeast growth rescue with the addition of exogenous ergosterol. The experimental data was complemented with docking simulations.

## RESULTS

### Selection of the best CgHMGR inhibitors

An evaluation was made of the possible antifungal activity of the thirteen compounds of series 1 and 2 and the seven compounds of series 3. The controls were the DMSO solvent and two compounds (α-asarone and fluconazole, at different concentrations) that reduce the synthesis of ergosterol in *C. glabrata* (Supplementary Figure 1) (Figure 1). The best inhibition of the growth of *C. glabrata* in solid YPD medium was exhibited by derivative **1c** (of series 1 and 2), and the substituted pyrrole derivate **5b** (of series 3) (Supplementary Figure 1) (Figure 1).

**Figure 1.**
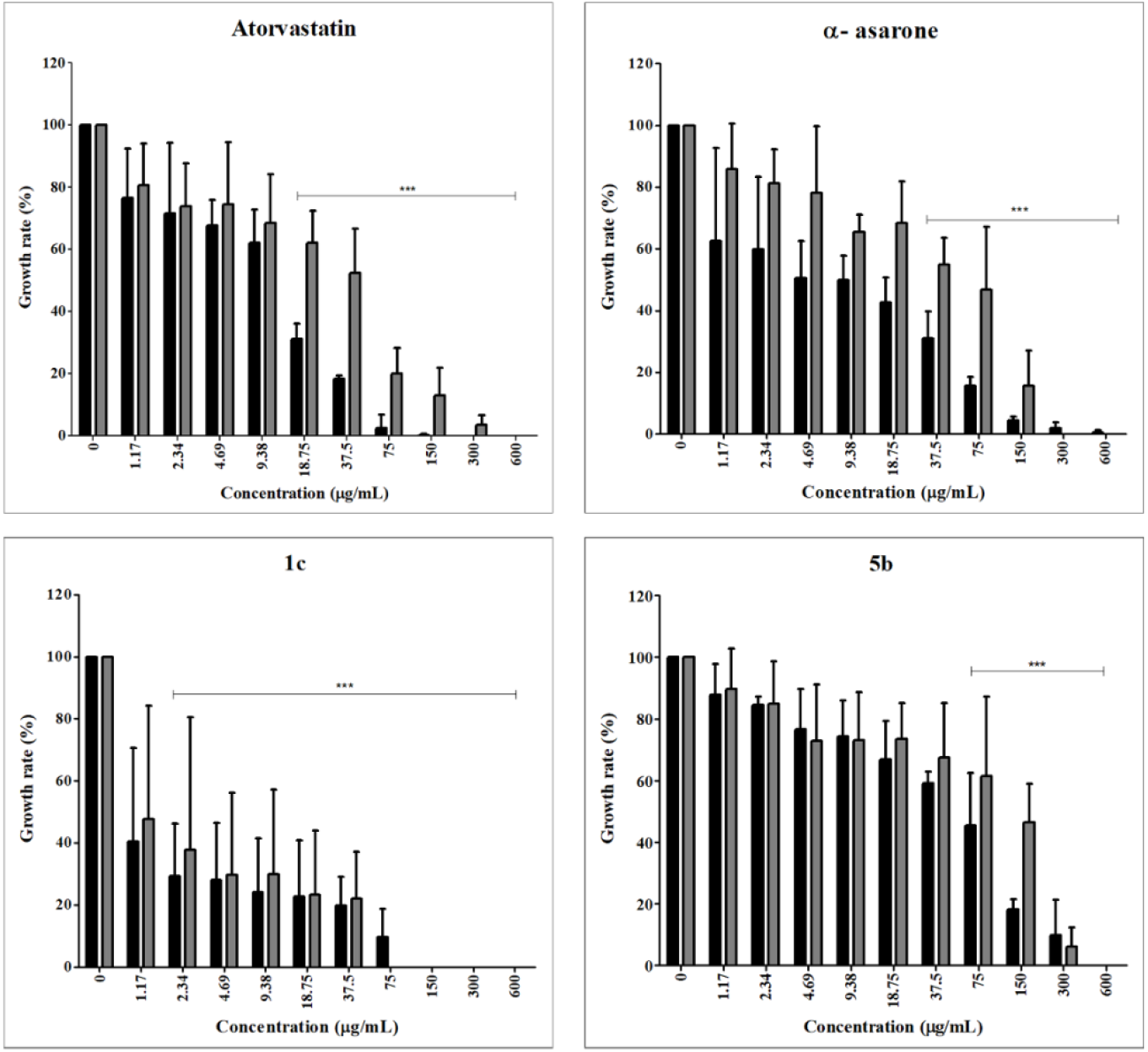
Inhibition of the growth of *Candida glabrata* CBS 138 (black bars) and *C. glabrata* 43 (gray bars) by HMGR inhibitors (antifungal reference and test compounds). As a control, the strains were grown without any inhibitor. The optical density was determined in a Thermo Scientific™ Multiskan™ FC microplate photometer at 620 nm. Growth rate values are expressed as the average of three independent assays ± SD. Significant differences were analyzed by two-way ANOVA. ***P<0.001.

### The HMGR inhibitors affect the viability of *Candida glabrata*

The phenotype of the strains was verified, being *C. glabrata* CBS 138 and *C. glabrata* 43, susceptibility and resistant to fluconazole, respectively. Once this was established, an evaluation was made of the *in vitro* antifungal activity of **1c, 5b**, α-asarone (from which **1c** is structurally related), and atorvastatin (an HMGR inhibitor and from which **5b** is structurally related). Both test compounds (**1c** and **5b**) and reference compounds (α-asarone and atorvastatin) were able to diminish the viability of the two strains of *C. glabrata*. **1c** at 75 µg/mL provided growth inhibition similar to atorvastatin and α-asarone at the same concentration, reducing yeast growth by up to 90% for the two strains. It was necessary to apply 300 µg/mL of **5b** to afford a similar percentage of inhibition (Figure 1) (Figure 2). As the concentration of the compound increased, the growth of the yeast strains decreased (Tables 1 and 2), indicating a dose-response effect. Compound **1c** presented lower IC_50_ and IC_70-90_ values than its control (α-asarone), **5b** and atorvastatin (Table 3).

**Table 1.**
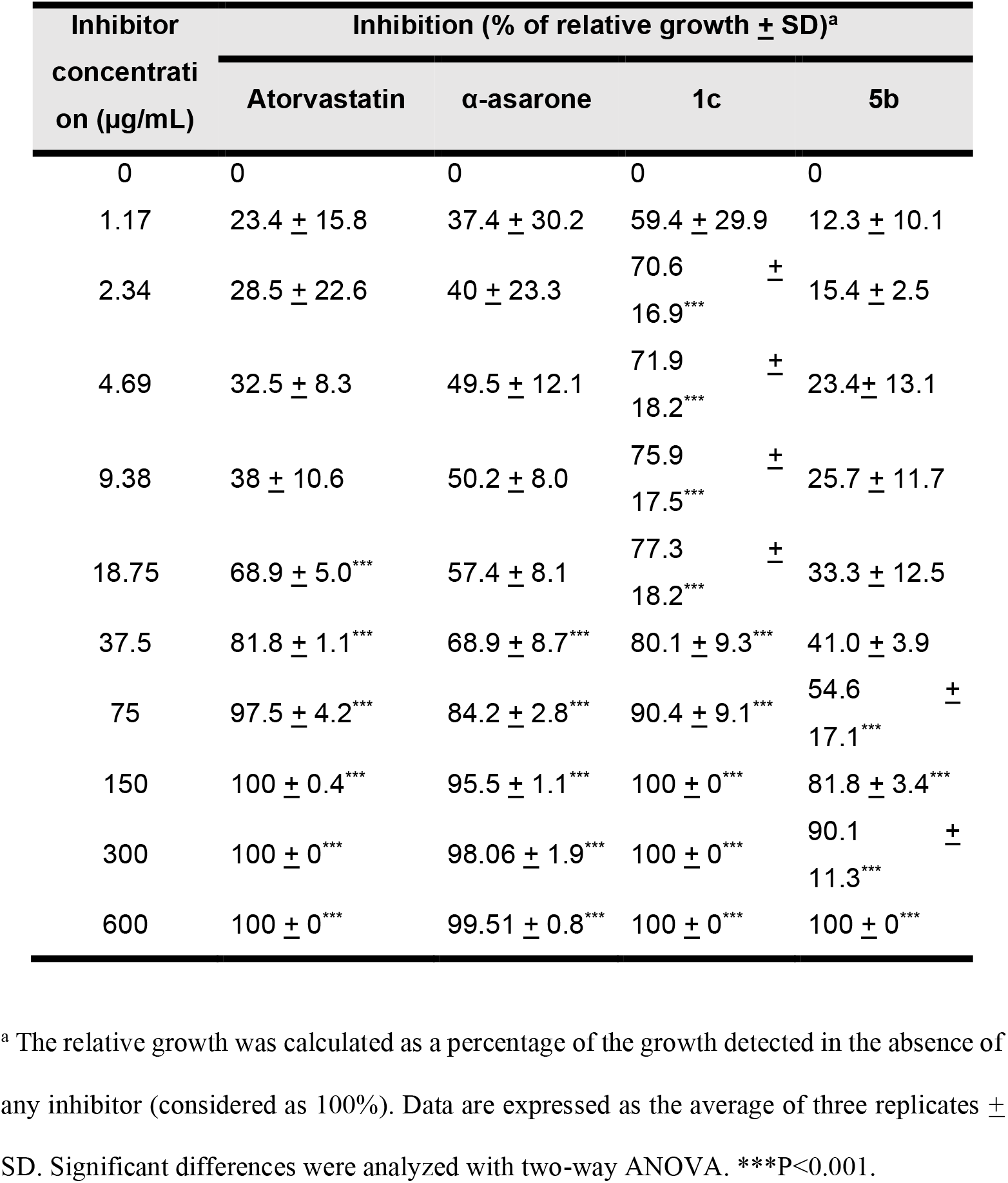
Effect of 1c, 5b, α-asarone and atorvastatin on the growth of *Candida glabrata* CBS 138.

| Inhibitor<br>concentration ($\mu\text{g/mL}$) | Inhibition (% of relative growth ± SD)<sup>a</sup> | | | |
| --- | --- | --- | --- | --- |
| | Atorvastatin | α-asarone | 1c | 5b |
| 0 | 0 | 0 | 0 | 0 |
| 1.17 | 23.4 ± 15.8 | 37.4 ± 30.2 | 59.4 ± 29.9 | 12.3 ± 10.1 |
| 2.34 | 28.5 ± 22.6 | 40 ± 23.3 | 70.6 ± 16.9*** | 15.4 ± 2.5 |
| 4.69 | 32.5 ± 8.3 | 49.5 ± 12.1 | 71.9 ± 18.2*** | 23.4 ± 13.1 |
| 9.38 | 38 ± 10.6 | 50.2 ± 8.0 | 75.9 ± 17.5*** | 25.7 ± 11.7 |
| 18.75 | 68.9 ± 5.0*** | 57.4 ± 8.1 | 77.3 ± 18.2*** | 33.3 ± 12.5 |
| 37.5 | 81.8 ± 1.1*** | 68.9 ± 8.7*** | 80.1 ± 9.3*** | 41.0 ± 3.9 |
| 75 | 97.5 ± 4.2*** | 84.2 ± 2.8*** | 90.4 ± 9.1*** | 54.6 ± 17.1*** |
| 150 | 100 ± 0.4*** | 95.5 ± 1.1*** | 100 ± 0*** | 81.8 ± 3.4*** |
| 300 | 100 ± 0*** | 98.06 ± 1.9*** | 100 ± 0*** | 90.1 ± 11.3*** |
| 600 | 100 ± 0*** | 99.51 ± 0.8*** | 100 ± 0*** | 100 ± 0*** |

<sup>a</sup> The relative growth was calculated as a percentage of the growth detected in the absence of any inhibitor (considered as 100%). Data are expressed as the average of three replicates ± SD. Significant differences were analyzed with two-way ANOVA. \*\*\*P<0.001.

**Table 2.** Effect of 1c, 5b, α-asarone and atorvastatin on the growth of *Candida glabrata* 43.

| Inhibitor<br>concentration<br>($\mu\text{g/mL}$) | Inhibition (% of relative growth ± SD)<sup>a</sup> | | | |
| --- | --- | --- | --- | --- |
| | Atorvastatin | α-asarone | 1c | 5b |
| 0 | 0 | 0 | 0 | 0 |
| 1.17 | 19.4 ± 13.4 | 14.2 ± 14.7 | 52.2 ± 36.5 | 10.7 ± 13.1 |
| 2.34 | 26.2 ± 13.7 | 18.9 ± 11.2 | 62.2 ± 42.7*** | 15.1 ± 13.7 |
| 4.69 | 25.7 ± 20.1 | 21.9 ± 21.6 | 70.3 ± 26.5*** | 27.2 ± 18.3 |
| 9.38 | 31.7 ± 15.8 | 34.6 ± 5.7 | 70.0 ± 27.1*** | 26.9 ± 15.5 |
| 18.75 | 38.0 ± 10.2*** | 31.7 ± 13.5 | 76.6 ± 20.7*** | 26.6 ± 11.7 |
| 37.5 | 47.8 ± 14.4*** | 45.1 ± 8.72*** | 77.9 ± 15.0*** | 32.6 ± 17.7 |
| 75 | 80.1 ± 8.3*** | 53.1 ± 20.3*** | 100 ± 0*** | 38.5 ± 25.7*** |
| 150 | 87.2 ± 8.9*** | 84.3 ± 11.5*** | 100 ± 0*** | 53.7 ± 12.6*** |
| 300 | 96.5 ± 3.0*** | 100 ± 0*** | 100 ± 0*** | 93.9 ± 6.2*** |
| 600 | 100 ± 0*** | 100 ± 0*** | 100 ± 0*** | 100 ± 0*** |

<sup>a</sup> The relative growth was calculated as a percentage of the growth detected in the absence of any inhibitor (considered as 100%). Data are expressed as the average of three replicates ± SD. Significant differences were analyzed with two-way ANOVA. \*\*\*P<0.001.

**Table 3.** MIC_50_ and MIC_70-90_ values of 1c, 5b, α-asarone and atorvastatin against *Candida glabrata*. The control consisted of the yeast strain cultivated without any inhibitor.

| Inhibitor | *C. glabrata* CBS 138 |  | *C. glabrata* 43 |  |
| --- | --- | --- | --- | --- |
| | MIC<sub>50</sub> (μg/mL) | MIC<sub>70-90</sub> (μg/mL) | MIC<sub>50</sub> (μg/mL) | MIC<sub>70-90</sub> (μg/mL) |
| Control | - | - | - | - |
| Atorvastatin | 13 | 37.5 | 40.1 | 195.2 |
| α-asarone | 9.38 | 113.5 | 60.5 | 204.5 |
| 1c | <1.17 | 75 | <1.17 | 58 |
| 5b | 62.3 | 300 | 131.7 | 108.2 |

**Figure 2.**
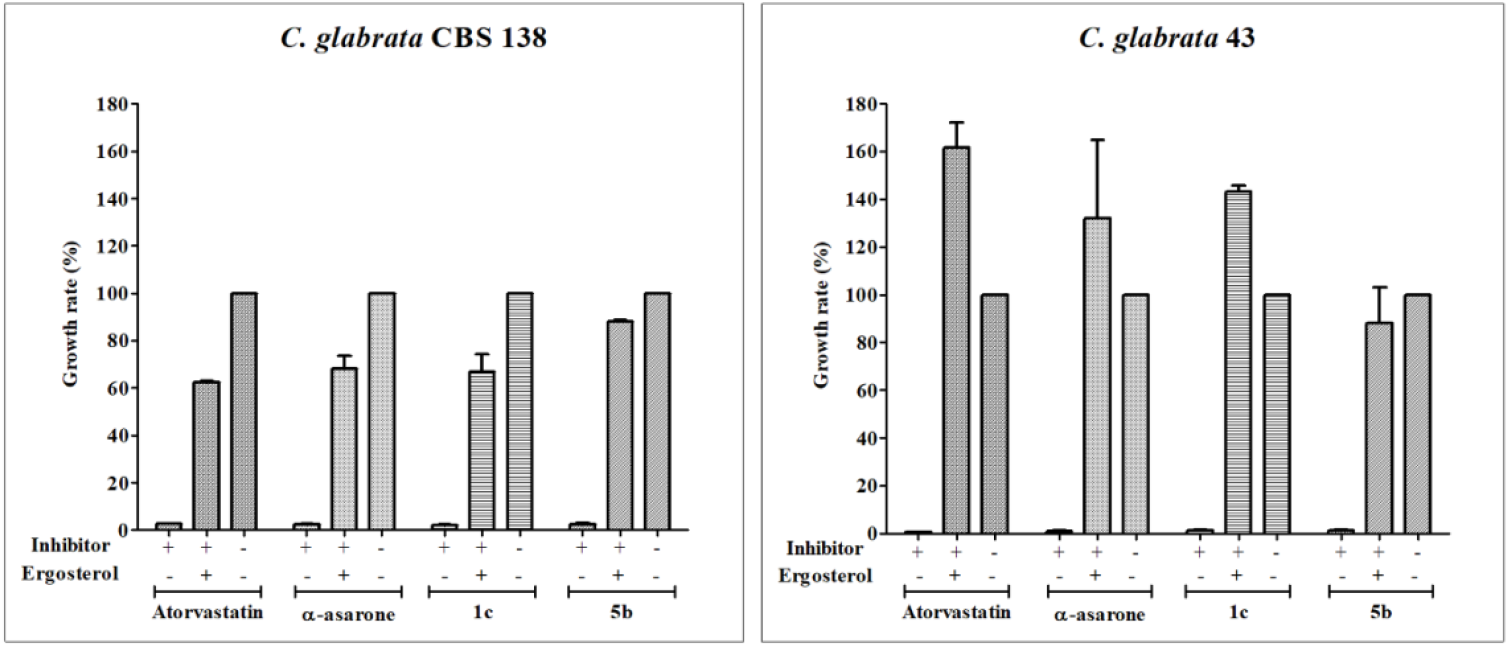
Yeast growth rescue experiment of *Candida glabrata* with ergosterol. After yeast growth was stopped by treatment with HMGR inhibitors (antifungal reference compounds and test compounds used at their IC_70-90_), the addition of exogenous ergosterol led to a recovery of the growth of *C. glabrata* CBS 138 and *C. glabrata* 43. The strains were also grown without any inhibitor as a control (considered as 100% growth). + represents addition of the inhibitor or ergosterol to the medium; – indicates the absence of the same. The optical density was determined in a Thermo Scientific™ Multiskan™ FC microplate photometer at 620 nm. Growth rate values (As_600_) are expressed as the average of three independent assays ± SD. ***P<0.001 compared to the assay without any inhibitor, based on the Student’s *t*-test.

### For *Candida glabrata* treated with inhibitors, growth recovered after adding ergosterol

A yeast growth rescue experiment was carried out to verify that the inhibition of the HMGR enzyme affects the levels of ergosterol, the final product of the biosynthesis pathway (Figure 2). The compounds were applied at the sublethal concentrations estimated in the previous experiment (MIC_70-90_). When exogenous ergosterol was subsequently added to the culture medium, yeast growth did indeed occur, in contrast to the lack of growth caused by the inhibitor. In some cases, such as with compound **1c** applied to *C. glabrata* 43, the recovery of yeast growth reached an even higher level than the control (the yeast cultured in the absence of an inhibitor). Thus, this finding confirmed that the compound derived from α-asarone altered the pathway for the production of ergosterol in *C. glabrata*, and more specifically that it targeted the synthesis of the HMGR enzyme.

### The test compounds (CgHMGR inhibitors) affect ergosterol biosynthesis in *Candida glabrata*

To explore the possible association between the loss of viability of *C. glabrata* and the inhibition of the production of ergosterol, the level of ergosterol in the yeasts was measured after 18 h of treatment with **1c, 5b**, simvastatin or α-asarone (the latter two as reference compounds; data not shown). The corresponding absorption spectra (Figure 3) contained the characteristic four peaks of sterols. The test compounds caused a reduction in the level of sterols in both the fluconazole-susceptibility and -resistant strains of *C. glabrata*.

**Figure 3.**
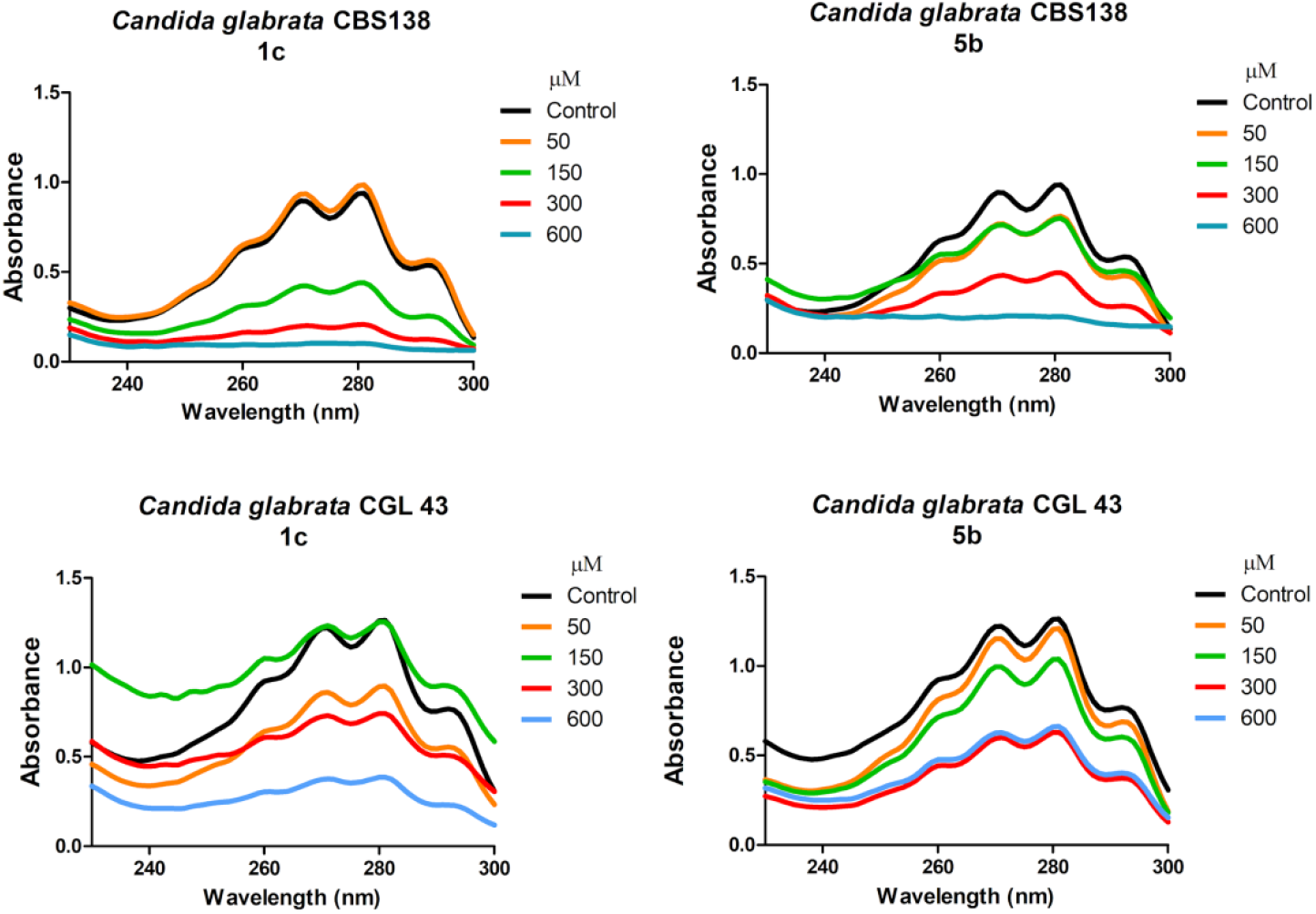
CgHMGR inhibitors **1c** and **5b** lowered the level of ergosterol. *C. glabrata* CBS138 and *C. glabrata* 43 were grown in YPD medium and treated with different concentrations (50, 100, 300 and 600 μM) of the inhibitors. The control was the YPD medium without any inhibitor or treated with the vehicle (DMSO) only. By spectrophotometrically scanning (from 230-300 nm) the extracted sterols (in the n-heptane layer), the presence or absence of ergosterol could be detected, as well as a possible reduction in the level of this sterol.

The absorption peak corresponding to 281.5 nm was used to quantify the concentration of ergosterol, allowing for the calculation of the percentage of inhibition of its synthesis (Table 4). In general, residual ergosterol levels were higher in the *C. glabrata* 43 versus *C. glabrata* CBS 138 strain. In both strains, a greater decrease in ergosterol was caused by **1c** than **5b**. Simvastatin and α-asarone served as positive controls for the inhibition of CgHMGR, since previous studies demonstrated their capability of inhibiting the recombinant HMGR of *C. glabrata* [8]. It is observed that the higher the concentration of the inhibitor, the greater the percentage of inhibition of ergosterol synthesis (Table 4).

**Table 4.** Percentage of ergosterol inhibition of *Candida glabrata* cells treated with HMGR enzyme inhibitors.

| Inhibitor | Concentration (μM) | *C. glabrata* CBS 138 | *C. glabrata* 43 |
| --- | --- | --- | --- |
| Control (W/I) | - | 100 | 100 |
| DMS Control | - | 100 | 100 |
| Simvastatin | 50 | 62.3 | 82.5 |
|  | 150 | 19.6 | 79.9 |
|  | 300 | 8.4 | 67.7 |
|  | 600 | 7.9 | 54.8 |
| α-asarona | 50 | 65.2 | 81.1 |
|  | 150 | 36.3 | 60.4 |
|  | 300 | 15.23 | 53.3 |
|  | 600 | 0.00 | 23.5 |
| 1c | 50 | 100.0 | 68.0 |
|  | 150 | 40.0 | 73.2 |
|  | 300 | 13.2 | 44.3 |
|  | 600 | 2.3 | 21.1 |
| 5b | 50 | 75.6 | 100.0 |
|  | 150 | 67.6 | 89.6 |
|  | 300 | 34.9 | 50.9 |
|  | 600 | 5.1 | 51.5 |

The level of ergosterol was calculated based on the absorbance obtained at 281.5 nm, expressing it as a percentage of the wet weight of the cells, as described by Arthington-Skaggs *et al*. [15]. *C. glabrata* was grown in YPD medium treated with different concentrations (50, 150, 300 and 600 μM) of the inhibitors: simvastatin, α-asarone, **1c** and **5b**. For the controls, the yeast was grown in YPD medium without any treatment or with DMSO only. The data represent the average of the three independent assays for each treatment. The previous results allowed for the calculation of the IC_50_, the concentration of the inhibitor that causes 50% inhibition of ergosterol synthesis in *C. glabrata* (Supplementary Table 1). **1c** had lower IC_50_ values than **5b** for both the *C. glabrata* CBS 138 (125 and 230 µM, respectively) and *C. glabrata* 43 (260 and >600 µM, respectively) strains.

### Docking suggests the interaction of the test compounds with HMGR of *Candida glabrata*

Docking simulations displayed the hypothetical interaction of the compounds with CgHMGR. The related values for **1c** and **5b** are shown in Table 5. **1c** has the highest binding energy *in silic*o, which correlates with the *in vitro* results (Table 1). Atorvastatin had the lowest binding energy (Table 5). The interaction of compounds **1c** and **5b** with the amino acid residues in the catalytic site is depicted in Figure 4. **1c** exhibited hydrogen bonds with a length of 2.58-2.99 Å between the hydroxyl groups at C-5 and C-8 and Glu93 and Asn192, respectively, as well as an electrostatic interaction of the O11 methoxy group with Met191. For **5b**, there were hydrogen bonds 2.19 and 19.7 Å in length between the hydroxyl group at C-5 and Met191, and between the carboxyl group at C-7 and Asp303, respectively. The interaction between atorvastatin and the HMGR catalytic site revealed that van der Waals interactions are predominant, although two hydrogen bonds are detected (19.7 and 22.7 Å) between the carboxyl group at C-17 and Gly341. Additionally, Asp303 interacted by hydrogen bonds with the carboxyl group at C-17 and the hydroxyl group at C-15 (Figure 4). The calculated binding energies of **1c** and **5b** (−5.99 and −5.71 kcal/mol, respectively) were better than those found for α-asarone and atorvastatin (4.53 and −2.13 kcal/mol, respectively) (Table 5).

**Table 5.**
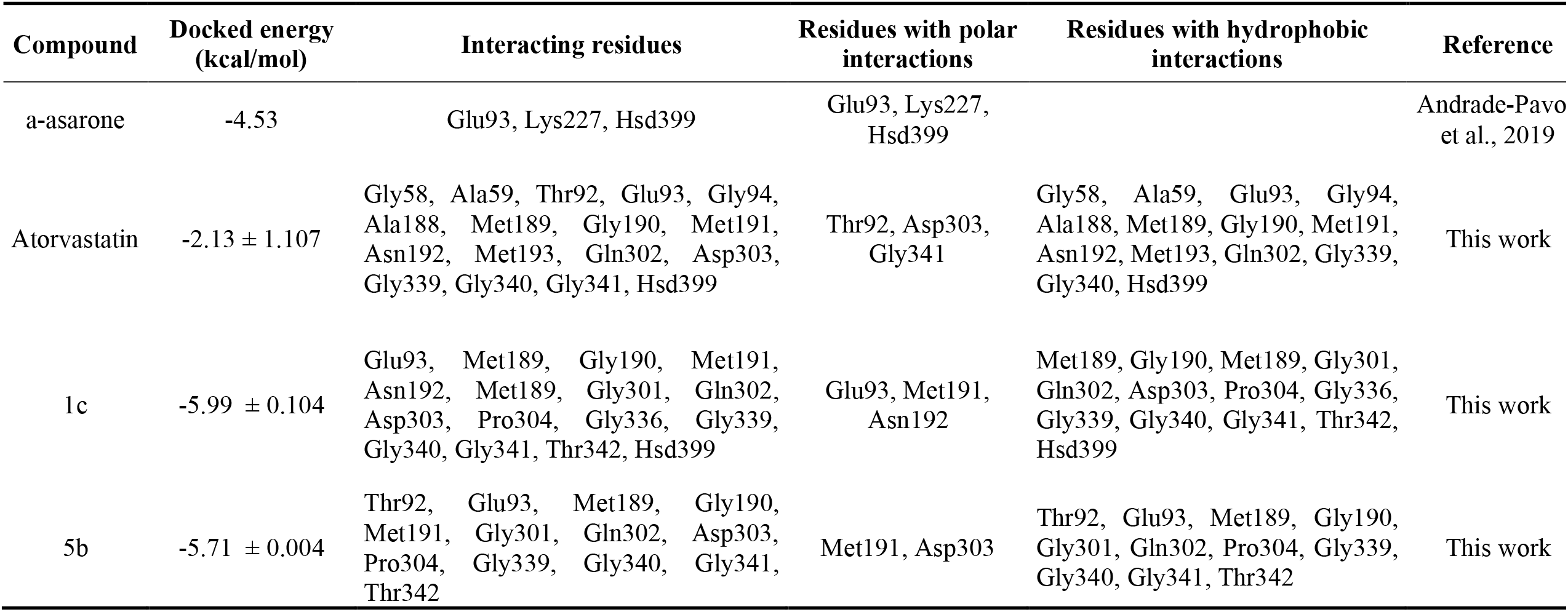
Docking data results of the binding mode between atorvastatin, 1c and 5b compounds on the catalytic site of CgHMGR.

**Figure 4.**
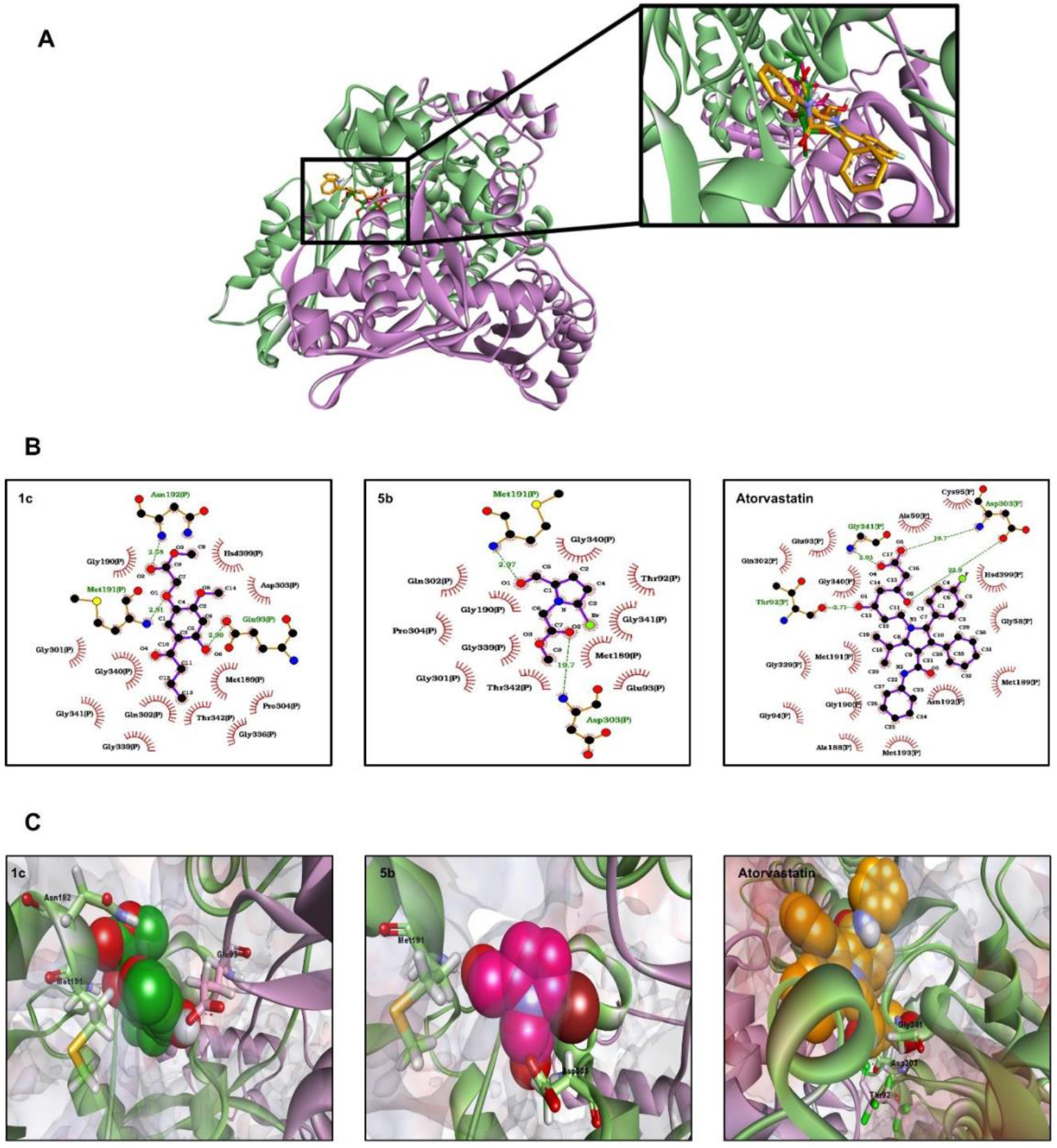
Schematic binding mode of **1c, 5b** and **atorvastatin** with the catalytic portion of CgHMGR. A) Structural model of CgHMGR, where the subunit a and b conforming to the catalytic domain are colored in green and purple respectively. A magnified visualization of the ligands interacting with the active site is shown in the black box. B) Predicted binding mode of **1c, 5b** and **atorvastatin** with the catalytic portion of CgHMGR. A docking simulation was conducted with AUTODOCK 4. In the 2D model obtained by using software, electrostatic and van der Waals interactions between the amino acid residues and the compounds are portrayed as red semicircles with rays. Hydrogen bonds are depicted by green dotted lines, and their size is denoted in Angstroms. C) 3D representation of the docking complexes between CgHMGR and **1c, 5b**, and **atorvastatin**. The α-helix and β-strand structures are represented as ribbons, colored in blue (subunit a) and purple (subunit b). The molecular surface electrostatic charges are shown. Ligands are represented as spheres. Amino acid residues that interact with ligands through H-bonds are represented as sticks. The figure is an original creation designed by Ortiz-Álvarez J. (co-author of this work) performed with the Discovery Studio 2020 Client and LigProt+ software.

## DISCUSSION

The problem of drug-resistant strains will always exist due to the process of natural evolution and selection of yeasts and bacteria [25]. Therefore, the probability of applying an effective treatment to patients would be increased by having a broad battery of antifungal agents from which to choose as well as distinct molecular targets among such drugs.

The HMGR enzyme (particularly CgHMGR) has for some time been proposed as a possible target, leading to the study of some cholesterol-lowering drugs (e.g., simvastatin and atorvastatin) as inhibitors of the growth of pathogenic yeasts [10,26]. According to *in vitro* evolutionary experiments, treatment of *C. glabrata* with some statins may allow for the selection of mutants. However, gene sequencing has not detected any changes in the catalytic domain of CgHMGR, indicating no effect on HMGR activity. *C. glabrata* is a useful model for examining resistance to statins and the precise molecular mechanisms of resistance to compounds that inhibit the CgHMGR enzyme [5].

In the current effort, three series of compounds were evaluated as inhibitors of *C. glabrata*. Two of the best derivatives were selected to determine their effect on yeast growth and ergosterol synthesis. Complementary studies were carried out with yeast growth rescue assays and docking simulations.

The compounds presently investigated were originally designed as lipid-lowering [11] and anti-inflammatory agents [12]. Their chemical structure could plausibly enable them to inhibit the activity of the CgHMGR enzyme. In fact, substituted pyrroles have been considered as antifungals [13,23] and their fungicidal activity is reported. However, the possible molecular target has not been previously explored in an in-depth manner.

Compounds such as statins (e.g., simvastatin and atorvastatin) and fibrates that inhibit HMGR have been administered to lower cholesterol levels in humans [27]. Additionally, they have been assessed as growth inhibitors of *Candida* spp., *Aspergillus* spp. and *Ustilago maydis* [7-10,14,26]. Based on its hypercholesterolemic activity, α-asarone underwent initial studies [27,28] that resulted in a finding of high toxicity. Thus, new derivative compounds have been designed and synthesized, and these have produced good activity against different fungi, such as *C. glabrata* and *Ustilago maydis* [9,14].

When the test compounds were examined *in vitro*, the growth inhibition of both strains of *C. glabrata* was better for **1c** than **5b** and α-asarone. On the other hand, **5b** did not induce a greater growth inhibition than its reference compound, atorvastatin. The latter statin, bearing a substituted pyrrolic ring, has already been proposed as an antifungal agent to inhibit the growth of *Candida* spp. [26]. Although the antifungal activity of **1c** has already been studied [11], this is the first evaluation, to our knowledge, of its effect on an opportunistic pathogenic yeast. Furthermore, the current investigation constitutes the first in-depth exploration of the mechanism of action and molecular target of the inhibitors.

According to the yeast growth rescue experiment, the test compounds likely inhibited the pathway for sterol biosynthesis [9,26]. The addition of ergosterol to *C. glabrata* CBS 138 resulted in a recovery of growth at a level below that of the control (without treatment with an inhibitor), while its addition to *C. glabrata* 43 led to growth that overcame the control level. This behavior can be explained by what is observed in the fluconazole*-*resistant *C. glabrata* strains, in which the consumption and metabolism of sterols might be affected by mutations in the *ERG11* gene. Moreover, the exposure of susceptible *C. glabrata* strains to fluconazole (an inhibitor of ergosterol synthesis) causes a coordinated action between the consumption and production of ergosterol. Hence, the present test compounds probably inhibit the pathway for sterol biosynthesis, as fluconazole does [30,31].

Since **1c** and **5b** inhibited ergosterol synthesis, they may reduce the activity of the CgHMGR enzyme [26]. A better inhibition of the production of ergosterol was found for **1c** in both strains of *C. glabrata* compared to its control (α-asarone) and **5b**. Of these compounds, **1c** had the lowest IC_50_. Previous publications have documented the capability of simvastatin, α-asarone, and derivatives of the latter to inhibit recombinant CgHMGR [8,9].

A correlation has been detected in *C. albicans* strains between their sensitivity to azoles and their total ergosterol concentration [15]. Therefore, it was important to demonstrate that the test compounds were capable of inhibiting the synthesis of ergosterol in both strains of *C. glabrata* (the fluconazole-susceptibility and -resistant strains).

The experimental results from the assays on yeast growth inhibition and the inhibition of ergosterol synthesis were complemented by docking simulations based on molecular coupling between the test compounds and CgHMGR. The binding energy values calculated for **1c** and **5b** were congruent with the *in vitro* findings for these two compounds. **1c** exhibited the lowest binding energies and the best *in vitro* inhibition of yeast growth. Better binding to the active site of CgHMGR was displayed by **1c** and **5b** than α-asarone and its derivates, based on the calculated binding energies of the present study for the former two and reports in the literature for the latter [9]. This supports the *in vitro* results, in which **1c** and **5b** showed the greatest inhibition of yeast growth and of ergosterol synthesis.

The high binding energy determined from the docking of **1c** and **5b** into the active site of CgHMGR may stem from the addition of the ester and hydroxyl groups to the molecule, elements that do not exist in the structure of α-asarone. The hydroxyl group of **1c** might play a crucial role in the proper binding mode of the compounds with HMGR [28,29]. Perhaps the chemical structure also confers a strong binding mode, considering the generation of hydrogen bonds with a short distance between atoms. On the other hand, the unsuitable binding mode of atorvastatin with CgHMGR possibly owes itself to steric interference of the chemical structure with a proper approach to the catalytic site of HMGR, as well as to the longer distance of the hydrogen bonds observed in the atorvastatin-CgHMGR complex (19.7 and 22.9 Å), which would confer weaker binding. Actually, atorvastatin has exhibited weak binding energy (−2.89 kcal/mol) with the catalytic site of human HMGR [32], substantiating the results obtained in this work with atorvastatin and CgHMGR. Interestingly, α-asarone, simvastatin and the substrate HMG-CoA presented almost identical high binding energy for the catalytic site of human HMGR [29]. Hence, the structural differences between human HMGR and CgHMGR may influence the binding mode.

Molecular modeling of proteins is a useful analytical technique that in the future should allow for the characterization of mutants in the *CgHmgr* gene, a phenotype resistant to antifungal inhibitors of the HMGR enzyme. Such resistance could be explained by changes in the protein related to its tertiary structure or by the capacity of inhibitors to bind with the amino acids of the catalytic site, among other possibilities. Indeed, molecular modeling analysis and mutations in the *ERG11* gene, encoding for the enzyme 14-alpha-lanosterol demethylase (CYP51), have already been carried out with distinct *C. albicans* strains. Thus, a molecular explanation can be provided for the resistance or sensitivity of these strains to different azoles [33].

## CONCLUSIONS

Three series of plausible inhibitors of the CgHMGR enzyme were designed, synthesized and tested for the inhibition of yeast growth. The two best candidates, **1c** (structurally related to fibrates) and **5b** (structurally related to atorvastatin), were chosen for further experiments. When comparing the results of these two compounds, treatment with the former led to a greater inhibition of yeast growth and ergosterol synthesis. The fact that the target of **1c** is the pathway for the synthesis of ergosterol was demonstrated by the decrease it caused in the level of ergosterol as well as the posterior rescue of yeast viability by the addition of exogenous ergosterol. According to the docking analysis, the present test compounds displayed a better binding mode with CgHMGR than α-asarone and atorvastatin, supporting the experimental results.

There are many advantages to the rational design of antifungal compounds that are derived from known drugs (statins, fibrates, etc.), have a defined chemical structure, and are directed at a specific target. Their pharmacokinetics and pharmacodynamics can be inferred, suggesting potential redesign strategies to make them more specific, more potent and less toxic. Based on the molecular modeling analysis, a plausible interaction of the inhibitory compound with the target protein is visualized and analyzed, thus providing insights into the possible mechanisms of resistance of a yeast to an antifungal agent. Such resistance might be explained on the basis of changes in the tertiary structure of the protein or in the binding mode of inhibitors with their target. The fibrate-related compound, **1c**, herein proved to be a good candidate for further research on its antifungal activity. Modifications of the compound should be considered to achieve greater specificity and potency. The derivatives could then be examined with *in vivo* animal models at a therapeutic dose. Other important areas to be explored are its toxicity and the inhibition of the recombinant CgHMGR enzyme.

## MATERIALS AND METHODS

### Strains and culture media

*C. glabrata* CBS 138 and *C. glabrata* 43 are susceptibility and resistant to fluconazole, respectively [9]. They were employed to examine the antifungal effect and ergosterol inhibition produced by the current test compounds. *C. glabrata* CBS 138 was donated by Dr. Bernard Dujon of the Pasteur Institute, Paris. *C. glabrata, C. albicans* and *C. krusei* strains were stored at −70 °C in 50% (v/v) anhydrous glycerol (Sigma-Aldrich). They were recovered in yeast extract-peptone-dextrose (YPD) medium (1% yeast extract, 2% casein peptone, and 2% dextrose anhydrous powder; J.T. Baker) at 37 °C under orbital shaking at 120 rpm, to be used as inoculum in the assays. The RPMI-1640 medium (Sigma-Aldrich) was prepared in accordance with the standard procedures of the Clinical and Laboratory Standards Institute (CLSI). For growth rescue assays, stock solutions of the yeasts were elaborated at a final concentration of 2.5% (v/v) in a mixture of Tween 80 and ethanol (1:1) (Sigma-Aldrich).

### Evaluation of the growth inhibition of *C. albicans* and *C. glabrata*

To identify the compounds with the greatest potential antifungal activity, all the compounds in the three series were examined, together with three reference compounds (fluconazole, α-asarone and simvastatin), for their effect on the growth of two strains of *C. albicans* and two strains of *C. glabrata*. Yeast cells were cultured in slightly stirred YPD medium at 37 °C for 24 h, and later adjusted to a density of 0.5 (As_600_) to obtain a new inoculum. A stock solution, prepared with dimethyl sulfoxide (DMSO) and 10 mM of each of the inhibitors, was added (50 μL) in a Petri dish to afford a final inhibitor concentration of 50, 300 or 600 μM. Subsequently, YPD medium (25 mL) was added, and the mixture was slightly stirred until a homogenous solid was formed. The solidified media were inoculated with 20 μL of each of the *Candida* strains, previously adjusted, in a section of the Petri dish and incubated at 37 °C for 24 h [14]. Based on this procedure, two inhibitors were selected for further experiments, **1c** from the fibrate derivates and **5b** from the substituted pyrroles.

### In vitro activity of the synthetic compounds against *Candida spp*

The effect of **1c** and **5b** on the growth of *C. glabrata* CBS 138 and *C. glabrata* 43 was evaluated by using the CLSI M27-A3 microdilution method. Briefly, stock solutions of antifungal compounds were prepared, from which the experimental concentrations were obtained in RPMI-1640 medium (Sigma-Aldrich). Fluconazole, simvastatin, atorvastatin and α-asarone served as reference compounds for examining susceptibility. *C. albicans* ATCC 10231 and *C. krusei* ATCC 14423 were the controls for sensitivity and resistance, respectively. The synthetic compounds were dissolved in DMSO at the time they were placed on the microplates, followed by incubation for 24 h at 37 °C. The volume of the solvent was less than 10% of the total volume to avoid problems of inhibition by the solvent. Growth was quantified by optical density in a Thermo Scientific™ Multiskan™ FC microplate spectrophotometer at 620 nm. The values of yeast growth are expressed as the average of three independent assays.

#### *Candida glabrata* growth rescue

To verify that inhibitors affect yeast viability by inhibiting ergosterol synthesis, a growth rescue experiment was conducted. Growth was first stopped by subjecting yeasts to the sublethal concentration (IC_70-90_) of one of the inhibitors, determined by the CLSI M27-A3 protocol (see section 2.3), and then ergosterol was added. Briefly, to each well of 96-well microplates were added 100 µL of one of the antifungal solutions (2x) prepared in RPMI-1640 medium (Sigma-Aldrich), followed by 80 µL of a yeast suspension adjusted to 1-5 × 10^6^ UFC/mL and diluted 1:1000 with RPMI-1640 medium (Sigma-Aldrich). A stock solution of ergosterol was prepared by dissolving 11 µg/mL in Tween 80/ethanol, and 20 µL of this solution was added to each well. The controls were yeasts cultured without any inhibitor (growth control) and those with an inhibitor but without sterol (growth rescue control).

### Statistical analysis

Data are expressed as the mean of three replicates ± standard deviation (SD). Differences between groups were examined with two-way analysis of variance (ANOVA), with the Bonferroni correction, and a 95% confidence interval. Statistical analyses were performed and graphs constructed with GraphPad Prism 5.0. Statistical significance was considered at P<0.001.

### Ergosterol quantification

Total sterols were extracted with a slightly modified version of the methodology reported by Arthington-Skaggs *et al*. [15]. Briefly, *C. glabrata* yeasts were grown in YPD medium by incubation at 37 °C for 24 h under constant agitation at 200 rpm. The cell culture was prepared by adjusting it to a density of 0.3 (As_600_) in different flasks containing 5 mL of YPD medium, followed by the addition of DMSO solvent as the control (Sigma-Aldrich, USA) or one of the inhibitors (simvastatin, α-asarone, **1c** or **5b** at 50, 150, 300 or 600 μM). For each treatment, the yeasts were incubated at 37 °C for 18 h under constant shaking at 200 rpm. Cells were harvested by centrifugation and washed with distilled water. After establishing the net weight of the pellet, it was mixed with 3 mL of an alcoholic solution of potassium hydroxide (KOH) (25 g of KOH and 35 mL of distilled water, brought to 100 mL with absolute ethanol) in a vortex for 1 min to extract the sterols [14-16]. The cell suspensions were incubated at 85 °C for 1 h, and then the sterols were extracted with a mixture of 1 mL of sterile distilled water and 3 mL of *n*-heptane by vigorously mixing in a vortex for 3 min. The *n*-heptane layer was spectrophotometrically scanned between 230 and 300 nm (BioSpectrometer, Eppendorf). The presence of ergosterol (As_281_._5_ peak) and 24 (28) dihydroxy-ergosterol (24 (28) DHE), a late intermediate (As_230_ peak), can be appreciated by the characteristic four-peaked spectrum indicating sterol absorption. The technique is also capable of revealing a decrease in the level of ergosterol. The absence of detectable levels is evidenced by a flattening of the curve [14-16].

### Docking of the test compounds on CgHMGR

The hypothetical three-dimensional structure of CgHMGR was obtained by homology modeling with MODELLER 9.13 software [17], using the crystallographic structure of human HMGR as the template (PDB entry: 1DQ8). The quality of the resulting model was evaluated by determining the stereochemical restrictions with a Ramachandran plot constructed on Procheck v.3.5.4 [18]. The structure was energetically minimized and equilibrated through molecular dynamic simulations on the NAMD2 program [19], which were performed in 2,000,000 steps for a total run time of 1 ns. The three-dimensional structure of the ligands, obtained with the ChemSketch program (www.acdlabs.com), was subjected to energy optimization and minimization with AVOGADRO software [20]. Docking simulations were conducted on AUTODOCK 4 [21], employing the parameters established by Andrade-Pavón *et al*. [9]. Docking results were computed based on a total of 100 runs and 1,250,000,000 generations, analyzed in AutodockTools and visualized on LigProt+ software [22].

### Synthesis of the compounds tested as potential antifungal agents

The fibrate-based derivates **1a**-**c, 2a**-**c** and **3a**-**c**, and 1,2-dihydroquinolines **4a**-**d**, constituted the first two series of compounds [11] (Figure 5). The substituted pyrrole derivatives comprised the third series, being **5a-d** and **6b-d** [12] (Figure 6). The brominated pyrroles **5b, 5c** and **6b-d** were designed because of its similarity to some pyrrole-based marine alkaloids known to exert both antifungal and antibacterial activity [13, 23-24].

**Figure 5.**
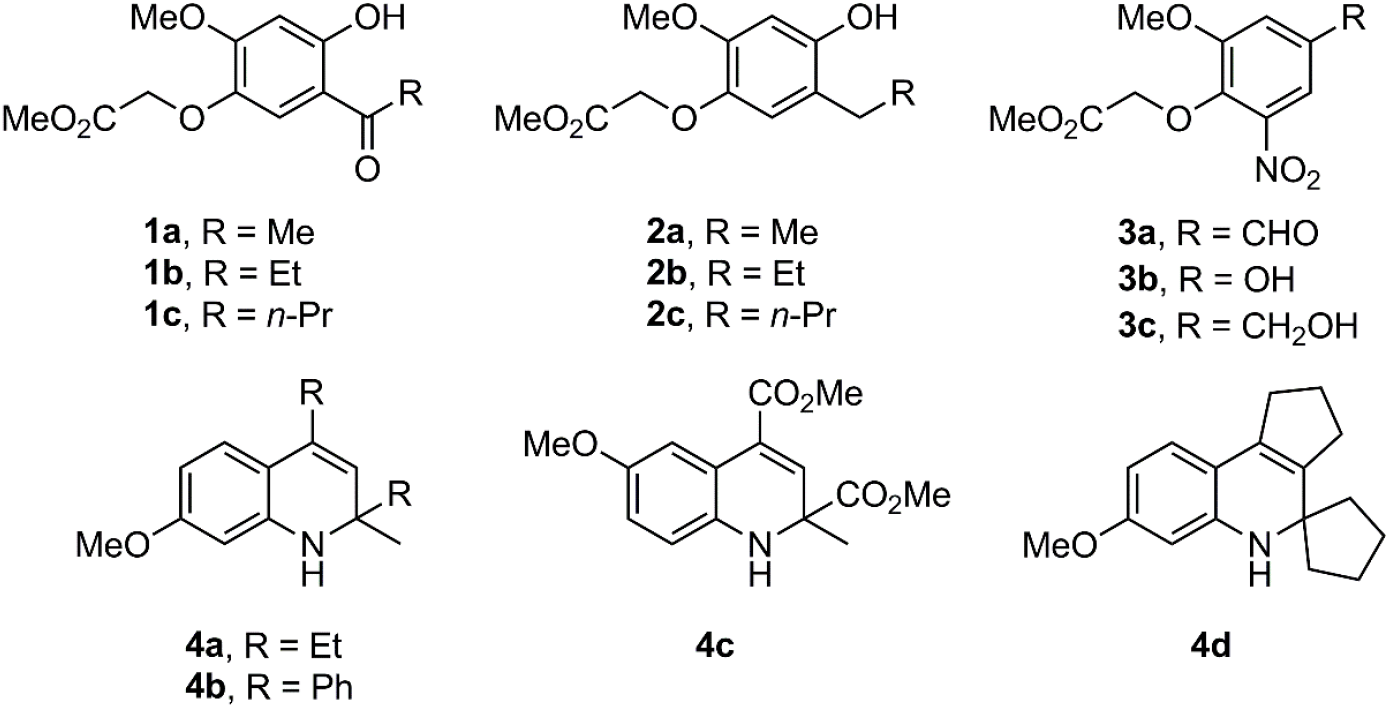
Structures of the fibrate-based analogues **1a-c, 2a-c** and **3a-c** (series 1), and 1,2-dihydroquinolines **4a-d** (series 2) [11].

**Figure 6.**
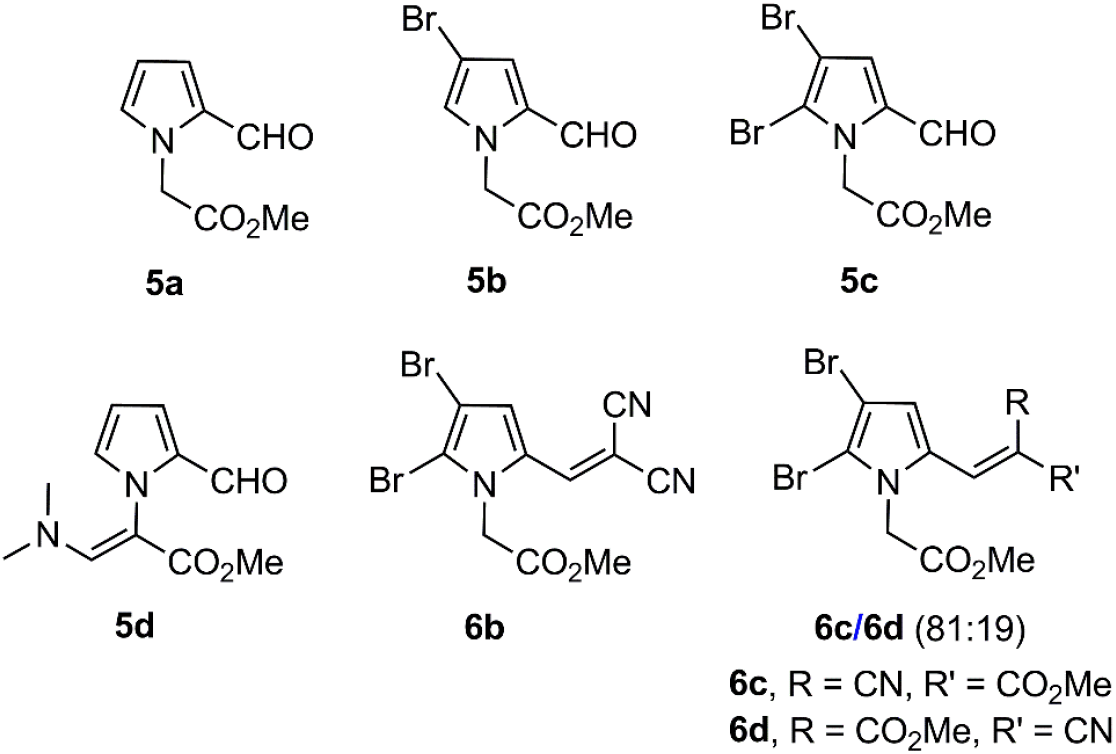
Structures of the substituted pyrroles **5a-d** and **6b-d** (series 3) [12].

### Synthesis of bromopyrroles 5b and 5c

The synthesis of **5a, 5d** and **6c-d** has been previously reported [11,12]. The preparation of bromopyrroles **5b** and **5c** was achieved by treatment of compound **5a** [12] with *N*-bromosuccinimide (NBS) as the brominating agent under mild reaction conditions (Scheme 1). Even though l.0 mol equivalent of NBS was employed, a mixture of bromopyrroles **5b** and **5c** was obtained. Due to the fact that they were easily separated by column chromatography, an excess of NBS (2.5 mol equiv.) was added to the reaction mixture to give **5b** and **5c** in 32% and 58% yields, respectively.

**Scheme 1.**
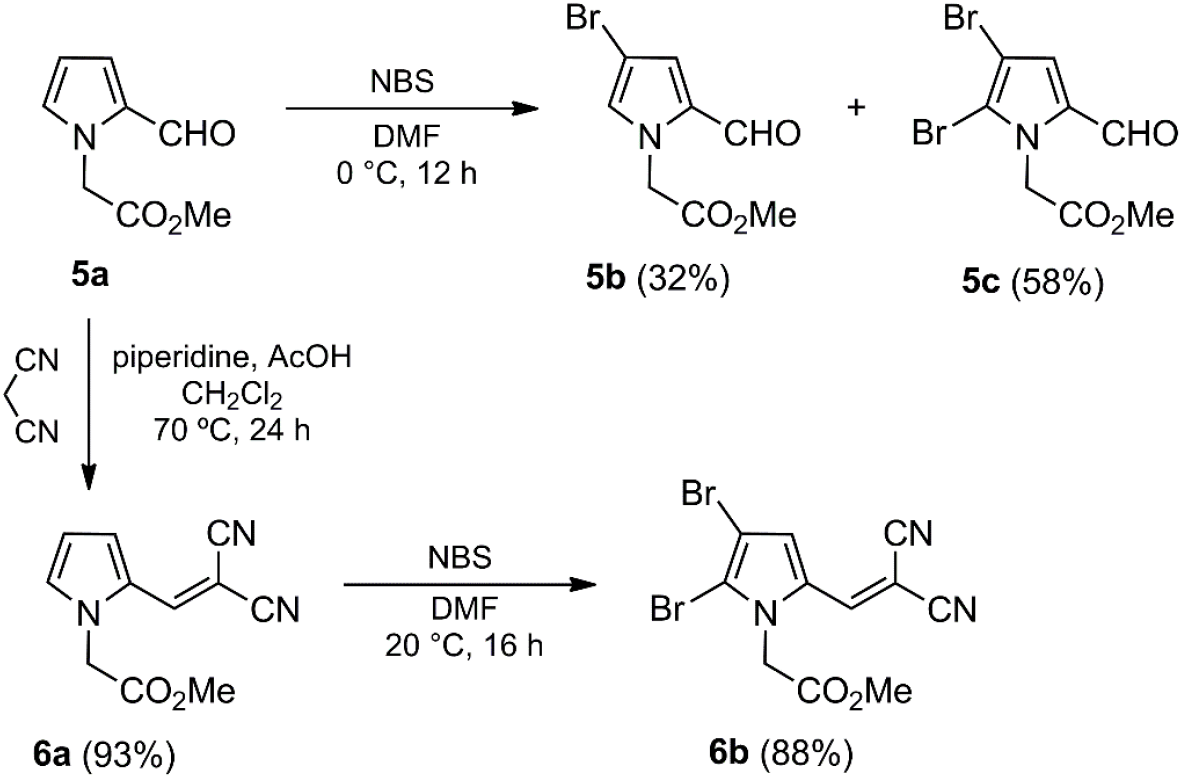
Synthesis of 4-bromopyrroles **5b, 5c** and **6b** from pyrrole **5a**.

The synthesis of dibromopyrrole **6b** was carried out by a two-step methodology. The first step consisted of a Knoevenagel reaction of **5a** with malononitrile under acid conditions [12] to provide **6a** in high yield (Scheme 1). Bromination of the latter with NBS (2.0 mol equiv.) in DMF, as the solvent, led to the desired product **6b** in good yield (88%) (Scheme 1).

### General information

Melting points were determined on a Krüss KSP 1N capillary melting point apparatus. IR spectra (ATR-FT or KBr) were recorded on a Perkin-Elmer 2000 spectrophotometer. ^1^H and ^13^C NMR spectra were captured on a Varian Mercury (300 MHz) instrument, with CDCl_3_ as the solvent and TMS as internal standard. Signal assignments were based on 2D NMR spectra (HMQC and HMBC). High-resolution mass spectra (HRMS) were obtained (in electron impact mode) on a Jeol JSM-GCMateII spectrometer. Analytical thin-layer chromatography was carried out using E. Merck silica gel 60 F254 coated 0.25 plates, visualized by using a long- and short-wavelength UV lamp. Flash column chromatography was conducted over Natland International Co. silica gel (230-400 and 230-400 mesh). All air moisture sensitive reactions were carried out under N_2_ using oven-dried glassware. CH_2_Cl_2_ and DMF (Sigma-Aldrich) were distilled over CaH_2_ (Sigma-Aldrich) prior to use. All other reagents (Sigma-Aldrich) were employed without further purification.

### Synthesis of bromopyrroles 5b and 5c

*Methyl 2-(4-bromo-2-formyl-1H-pyrrol-1-yl)acetate* (**5b**). *Methyl 2-(2,3-dibromo-5-formyl-1H-pyrrol-1-yl)acetate* (**5c**). To a stirring solution of **5a** (0.100 g, 0.60 mmol) in anhydrous DMF (5 mL) at 0 °C, a solution of NBS (0.267 g, 1.50 mmol) in anhydrous DMF (2 mL) was added dropwise, and the mixture stirred at 0 ° C for 12 h. A mixture of water/hexane/EtOAc (1:0.5:0.5) (20 mL) was added, the organic layer dried (Na_2_SO_4_) and the solvent removed under vacuum. The residue was purified by column chromatography over silica gel (30 g/g crude, hexane/EtOAc, 9:1) leading to **5b** (0.062 g, 32%) as a yellow solid and **5c** (0.112 g, 58%) as a yellow oil.

Data of **5b**: R*f* 0.43 (hexane/EtOAc, 7:3); mp 203-205 °C. IR (film): 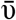 3121, 2954, 1754, 1666, 1392, 1365, 1219, 1092, 923, 771 cm^-1. 1^H NMR (300 MHz, CDCl_3_): δ 3.78 (s, 3H, CO_2_C*H*_3_), 5.03 (s, 2H, C*H*_2_), 6.92 (br dd, *J* = 1.8, 1.2 Hz, 1H, H-5’), 6.98 (d, *J* = 1.8 Hz, 1H, H-3’), 9.47 (d, *J* = 0.9 Hz, 1H, C*H*O). ^13^C NMR (75.4 MHz, CDCl_3_): δ 50.1 (*C*H_2_), 52.7 (CO_2_*C*H_3_), 97.5 (C-4’), 125.2 (C-3’), 131.2 (C-5’), 131.6 (C-2’), 168.2 (*C*O_2_CH_3_), 179.3 (*C*HO). HRMS (EI): *m/z* [M^+^] calcd for C_8_H_8_BrNO_3_: 244.9688; found: 244.9690.

Data of **5c**: R*f* 0.69 (hexane/EtOAc, 7:3); IR (film): 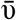 2955, 1755, 1668, 1450, 1397, 1363, 1218, 1005, 810, 776 cm^-1. 1^H NMR (300 MHz, CDCl_3_): δ 3.78 (s, 3H, CO_2_C*H*_3_), 5.25 (s, 2H, C*H*_2_), 7.05 (s, 1H, H-4’), 9.32 (s, 1H, C*H*O). ^13^C NMR (75.4 MHz, CDCl_3_): δ 49.0 (*C*H_2_), 52.7 (CO_2_*C*H_3_), 101.1 (C-3’), 118.6 (C-5’), 125.3 (C-4’), 132.4 (C-2’), 167.5 (*C*O_2_CH_3_), 178.2 (*C*HO). HRMS (EI): *m/z* [M^+^] calcd for C_8_H_7_Br_2_NO_3_: 322.8793; found: 322.8791.

### Synthesis of pyrroles 6a and 6b

*Methyl 2-(2-(2,2-dicyanovinyl)-1H-pyrrol-1-yl)acetate* (**6a**). In a threaded ACE glass pressure tube with a sealed Teflon screw cap and magnetic stirring bar, a solution of **5a** (0.100 g, 0.60 mmol), malononitrile (0.044 g, 0.66 mmol), piperidine (0.026 g, 0.30 mmol) and glacial AcOH (0.029 g, 0.48 mmol) in anhydrous CH_2_Cl_2_ (5 mL) was heated at 70 °C for 24 h. The reaction mixture was diluted with CH_2_Cl_2_ (50 mL) and washed with water (25 mL) and an aqueous saturated solution of NaHCO_3_ until neutral. The organic layer was dried (Na_2_SO_4_) and the solvent removed under vacuum. The residue was purified by column chromatography over silica gel (20 g/g crude, hexane/EtOAc, 9:1) to afford **6a** (0.12 g, 93%) as a yellow solid. R*f* 0.51 (hexane/EtOAc, 8:2); mp 203-205 °C. IR (KBr): 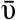 3132, 2992, 2220, 1751, 1583, 1476, 1399, 1350, 1328, 1239, 1169, 1132, 1088, 994, 758, 732 cm^-1. 1^H NMR (300 MHz, CDCl_3_): δ 3.83 (s, 3H, CO_2_C*H*_3_), 4.80 (s, 2H, C*H*_2_CO_2_Me), 6.49 (ddd, *J =* 4.5, 2.4, 0.6 Hz, 1H, H-4’), 7.10 (dd, *J* = 2.4, 1.5 Hz, 1H, H-5’), 7.38 (s, 1H, H-1”), 7.73 (ddd, *J* = 4.5, 1.5, 0.6 Hz, 1H, H-3’). ^13^C NMR (75.4 MHz, CDCl_3_): δ 48.3 (*C*H_2_CO_2_Me), 53.3 (CO_2_*C*H_3_), 72.5 (C-2”), 113.4 (C-4’), 114.0 (*C*N), 115.3 (*C*N), 121.1 (C-3’), 127.2 (C-2’), 131.6 (C-5’). 142.7 (C-1”), 167.3 (*C*O_2_CH_3_). HRMS (EI): *m/z* [M^+^] calcd for C_12_H_9_N_3_O_2_: 215.0695; found: 215.0694.

### Methyl 2-(2,3-dibromo-5-(2,2-dicyanovinyl)-1H-pyrrol-1-yl)acetate (6b)

To a stirring solution of **6a** (0.100 g, 0.47 mmol) in anhydrous DMF (3 mL) at 0 °C, a solution of NBS (0.166 g, 0.93 mmol) in anhydrous DMF (3 mL) was added dropwise, and the mixture stirred at 20 ° C for 16 h. A mixture of water/hexane/EtOAc (0.5:1:1) (30 mL) was added, the organic layer dried (Na_2_SO_4_) and the solvent removed under vacuum. The residue was purified by column chromatography over silica gel (20 g/g crude, hexane/EtOAc, 7:3) to give **6b** (0.25 g, 88%) as a yellow solid. R*f* 0.44 (hexane/EtOAc, 8:2); mp 267-269 °C. IR (KBr): 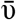 3004, 2956, 2223, 1741, 1583, 1420, 1391, 1339, 1243, 1167, 1125, 1004, 983, 815, 738, 687 cm^-1. 1^H NMR (300 MHz, CDCl_3_): δ 3.85 (s, 3H, CO_2_C*H*_3_), 4.89 (s, 2H, C*H*_2_CO_2_Me), 7.31 (d, *J =* 0.3 Hz, 1H, H-1”), 7.76 (d, *J =* 0.3 Hz, 1H, H-4’). ^13^C NMR (75.4 MHz, CDCl_3_): δ 47.8 (*C*H_2_CO_2_Me), 53.5 (CO_2_*C*H_3_), 75.1 (C-2”), 104.8 (C-3’), 113.3 (*C*N), 114.4 (*C*N), 118.6 (C-5’), 121.5 (C-4’), 128.3 (C-2’), 141.4 (C-1”), 166.1 (*C*O_2_CH_3_). HRMS (EI): *m/z* [M^+^] calcd for C_11_H_7_Br_2_N_3_O_2_: 370.8905; found: 370.8905.

### Reference compounds for the tests of the three series of potential antifungal compounds 1a-c, 2a-c, 3a-c, 4a-d, 5a-d and 6b-d

Depending on the experiment, different inhibitors served as the reference compounds. In the case of sensitivity tests and docking analysis, α-asarone was the control for the fibrate-based derivatives **1a-c, 2a-c, 3a-c** (series 1) and 1,2-dihydroquinolines **4a-d** (series 2) and atorvastatin for the substituted pyrroles **5a-d** and **6b-d** (series 3). In the experiment to determine the effect of the compounds on the biosynthesis of ergosterol, simvastatin and α-asarone were employed. It has been reported that these two compounds are capable of inhibiting recombinant Cg-HMGR, thus affecting the production of ergosterol [9].

## Acknowledgments

DMA, AGS, BRA, JOA, CBC and CHE appreciate graduate scholarship awarded by CONACyT as well as the scholarship complements furnished by the SIP-IPN (BEIFI). The authors would like to thank Bruce Allan Larsen for proofreading the manuscript. CH-R, GC-C, JT. and LV-T. are fellows of the *Estímulos al Desempeño de los Investigadores* (EDI)-IPN and *Comisión de Operación y Fomento de Actividades Académicas* (COFAA)-IPN programs.

## ABBREVIATIONS

ANOVA: analysis of variance
CgHMGR: 3-hydroxy-methyl-glutaryl-CoA reductase in *Candida glabrata*
CLSI: Clinical and Laboratory Standards Institute
DHE: dihydroxy-ergosterol
DMSO: dimethyl sulfoxide
HMGR: 3-hydroxy-methyl-glutaryl-CoA reductase
KOH: potassium hydroxide
NBS: *N*-bromosuccinimide
SD: standard deviation
YPD: yeast extract-peptone-dextrose medium

## SUPPLEMENTAL MATERIAL

**Supplementary Figure 1.**
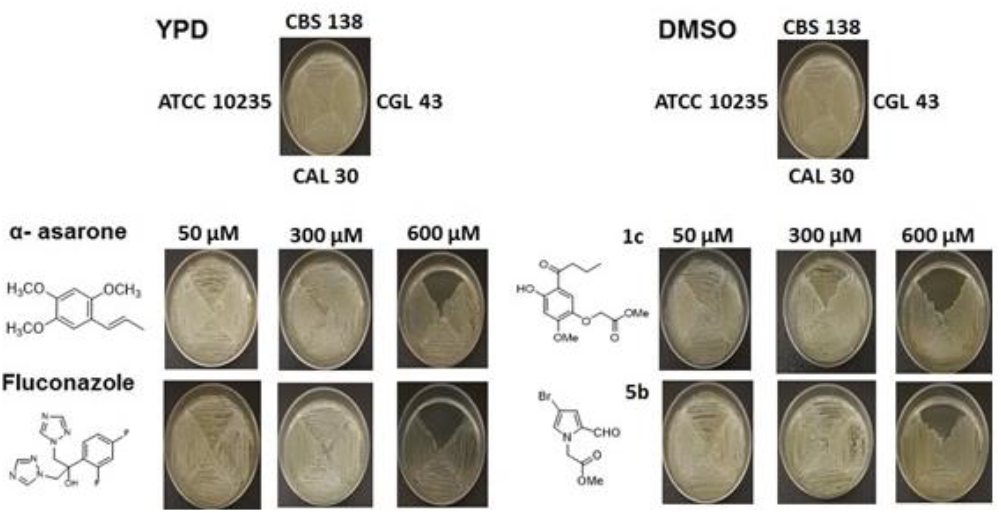
Effect of the inhibitors **1c** and **5b** on yeast growth. *C. albicans* (ATCC 10235, CAL30) and *C. glabrata* (CBS138, CGL43) were grown on solid YPD medium with 50, 300 and 600 μM of the inhibitors. The two *C. albicans* strains served as the controls. α-asarone and fluconazole were used as controls that inhibited ergosterol synthesis.

**Supplementary Table 1.**
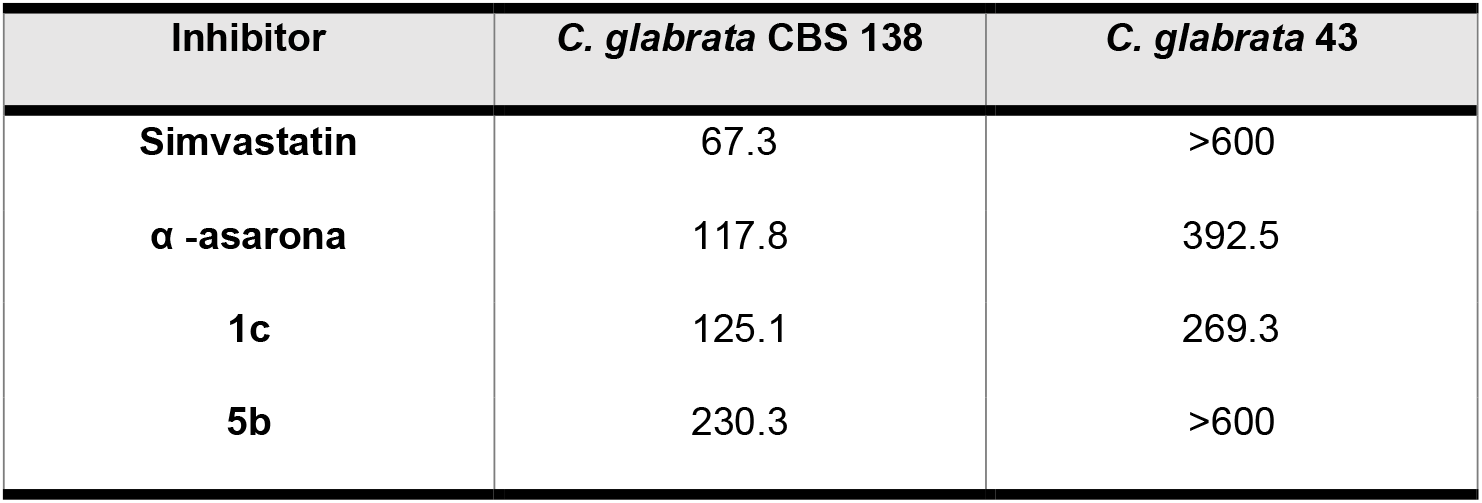
The IC_50_(µM) is given for each compound. This is the concentration that inhibits 50% of the ergosterol in *C. glabrata*.

